# Repair of CRISPR-guided RNA breaks enables site-specific RNA editing in human cells

**DOI:** 10.1101/2023.08.29.555404

**Authors:** Anna Nemudraia, Artem Nemudryi, Blake Wiedenheft

## Abstract

Genome editing with CRISPR RNA-guided endonucleases generates DNA breaks that are resolved by cellular DNA repair machinery. However, analogous methods to manipulate RNA remain unavailable. Here, we show that site-specific RNA breaks generated with RNA-targeting CRISPR complexes are repaired in human cells, and this repair can be used for programmable deletions in human transcripts that restore gene function. Collectively, this work establishes a technology for precise RNA manipulation with potential therapeutic applications.

**One-Sentence Summary:** CRISPR-guided RNA breaks are repaired in human cells, and this RNA repair can be used for programmable editing of human transcriptomes.

## Main Text

CRISPR RNA-guided endonucleases have enabled programable DNA cleavage and the development of new therapeutics (*1, 2*). The first generation of CRISPR genome editing used Cas9 nuclease to make double-stranded DNA breaks that are repaired by the cell, leading to edits at the location of the break (*3*). These technologies have enriched our mechanistic understanding of DNA repair, and new insights into repair are used to improve methods for genome editing (*4*).

While DNA editing has the potential to cure genetic diseases, it can result in unintended changes to the genome (*5*) and activate cellular stress responses that result in toxicity (*6, 7*). In contrast to DNA editing, RNA editing can be used to alter the cellular program without introducing heritable modifications in the DNA. However, options for editing RNA are limited. Cas13, a type VI CRISPR RNA-guided endoribonuclease, has been used to knock down cellular transcripts, but target recognition by Cas13 also activates collateral non-sequence-specific nuclease activity that degrades non-target RNAs (*8, 9*). Nuclease-inactivated mutants of Cas13 (dCas13) have been used for inducing exon skipping (*10*), trans-splicing of synthetic RNA payloads (*11*), or sequence-specific delivery of base-editing enzymes for adenosine-to-inosine or cytosine-to-uracil conversions (*12, 13*). However, programmable deletions in RNA, analogous to those introduced by DNA editors, have not been described.

To address the need for versatile and facile manipulation of RNA, we have developed a technology for sequence-specific RNA editing in living cells. This technology uses type III CRISPR complexes for programmable cleavage of target RNA. Unlike Cas13, which has non-sequence-specific nuclease activity, type III complexes exclusively cleave the target RNA in six nucleotide increments (*14, 15*). We repurpose this precise cleavage activity to excise six nucleotide segments from human transcripts and show that resulting RNA fragments are repaired in human cells, which produces programmable deletions in target RNA. We apply this method to delete premature stop codons in RNA and rescue protein expression, which demonstrates the potential of this technique for treating genetic diseases. Overall, this work establishes a technology for RNA manipulation in living cells and provides a model for discovering new RNA repair pathways.

### Programmable deletions in human transcripts

The programmable RNase activity of type III complexes has been previously used to knock down RNA targets in several cell types (*14, 16-19*), but we hypothesized that CRISPR-guided RNA breaks are repaired by cellular RNA repair machinery, which could be leveraged for RNA editing (**Fig. 1A**). To test this hypothesis, we transfected human cell cultures (i.e., 293T cells) with plasmids that express type III CRISPR-associated proteins from *Streptococcus thermophilus* (Csm2 – Csm5, Cas10, and Cas6) (*16*) fused to a nuclear localization signal (NLS) and RNA guides (SthCsm complex) complementary to sequences of *PPIB* or *PARK7* messenger RNAs (**fig. S1**).

**Fig. 1.**
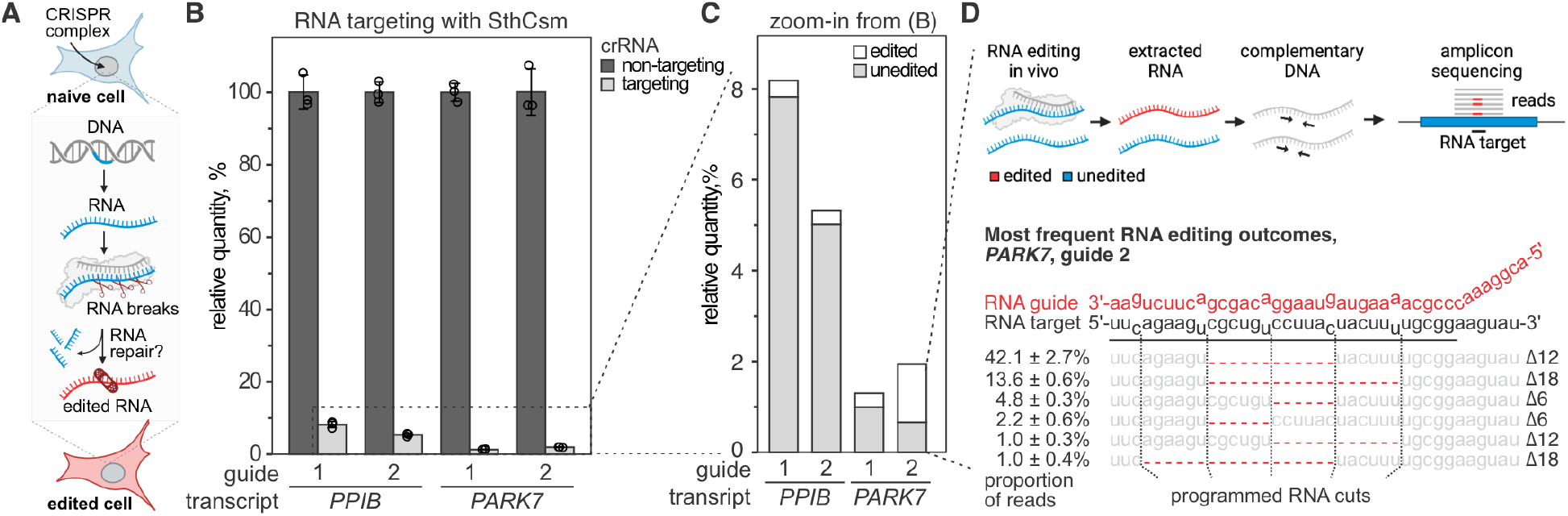
Programmable deletions of RNA with RNA-guided type III-A CRISPR complexes. **A**) Diagram of RNA editing in eukaryotic cells through sequence-specific RNA cleavage and RNA repair. **B**) Human cells (293T) were transfected with plasmids encoding for type III CRISPR complex of *Streptococcus thermophilus* (SthCsm), and RNA guides targeting *PPIB* or *PARK7* messenger RNAs. Target transcripts were quantified with RT-qPCR, and the qPCR signal was normalized to *ACTB* and non-targeting guide RNA control using the ΔΔCt method. Data is shown as mean ± one standard deviation of three biological replicates. **C**) Zoom-in from panel B. Deep sequencing was used to quantify the proportion of signal that is derived from edited RNA. **D**) *Top:* schematics of deep sequencing approach used to quantify RNA editing. Sequencing reads were aligned to the reference sequence, and modifications at the target site were quantified. *Bottom*: top five most frequent RNA editing outcomes in *PARK7* transcript (guide 2). Dotted lines indicate the positions of RNA breaks by the SthCsm complex. Red dashes depict deletions (Δ) identified in the sequencing data. The proportion of reads with deletions is shown as mean ± one standard deviation of three biological replicates.

To measure target transcript levels, we designed RT-qPCR primers that flank the targeted sequence (**fig. S1, C and F**). Expression of the SthCsm complex with RNA guides results in >90% reduction of targeted transcript levels compared to cells transfected with a non-complementary guide RNA (**Fig. 1B**), which is similar to the knockdown efficiencies previously reported for other guides (*16*). We hypothesized that the remaining target RNA was either not cleaved, cleaved and repaired without changing the original sequence, or cleaved and repaired with indels at the target site (**Fig. 1C**). To test this hypothesis, we deep-sequenced the *PPIB* and *PARK7* amplicons to determine target editing outcomes and efficiencies (**Fig. 1D**, top). In addition to the intact sequence, we detect 6, 12, 18, or 24 nucleotide deletions in each of the two RNA targets, which are not seen in controls transfected with a non-targeting guide and are consistent with the six-nucleotide cleavage pattern of the SthCsm complex (**Fig. 1D**, bottom). SthCsm-dependent target site deletions were detected in 4.4 ± 0.6% (guide 1) and 5.9 ± 0.6% (guide 2) of sequenced *PPIB* amplicons, and 23.9 ± 2.3% (guide 1) and 65.8 ± 3.6% (guide 2) of sequenced *PARK7* amplicons (**Fig. 1D, fig. S1**). This data demonstrates that most of the target RNA is degraded, but some of the RNA is repaired in human cells.

### RTCB ligase repairs type III CRISPR-mediated RNA breaks

RNA cleavage by type III CRISPR complexes leaves a 2’,3’-cyclic phosphate and a 5’-hydroxyl at each cut site (*14*). Mammalian RNA ligase RTCB joins 2′,3′-cyclic phosphate, and a 5′-hydroxyl to seal RNA breaks when introns are excised during tRNA splicing and unconventional splicing of *XBP1* mRNA in the unfolded protein response (*20*) (**Fig. 2A**). Therefore, we hypothesized that RTCB ligase repairs transcripts targeted by SthCsm cleavage in human cells. To test this hypothesis, we used Cas9 nuclease to knock out the *RTCB* gene in 293T (**fig. S2**). There are three copies of the *RTCB* gene in 293T cells, and after screening 41 clones with Sanger sequencing, we have not identified any with a complete knockout (*21*) (**fig. S2D**). These results are consistent with earlier work showing that *RTCB* is essential (*20*). However, we have identified clones with a knockout of two copies of the gene, which results in a substantial depletion of the RTCB protein (**Fig. 2B; fig. S2, E and F**).

**Fig. 2.**
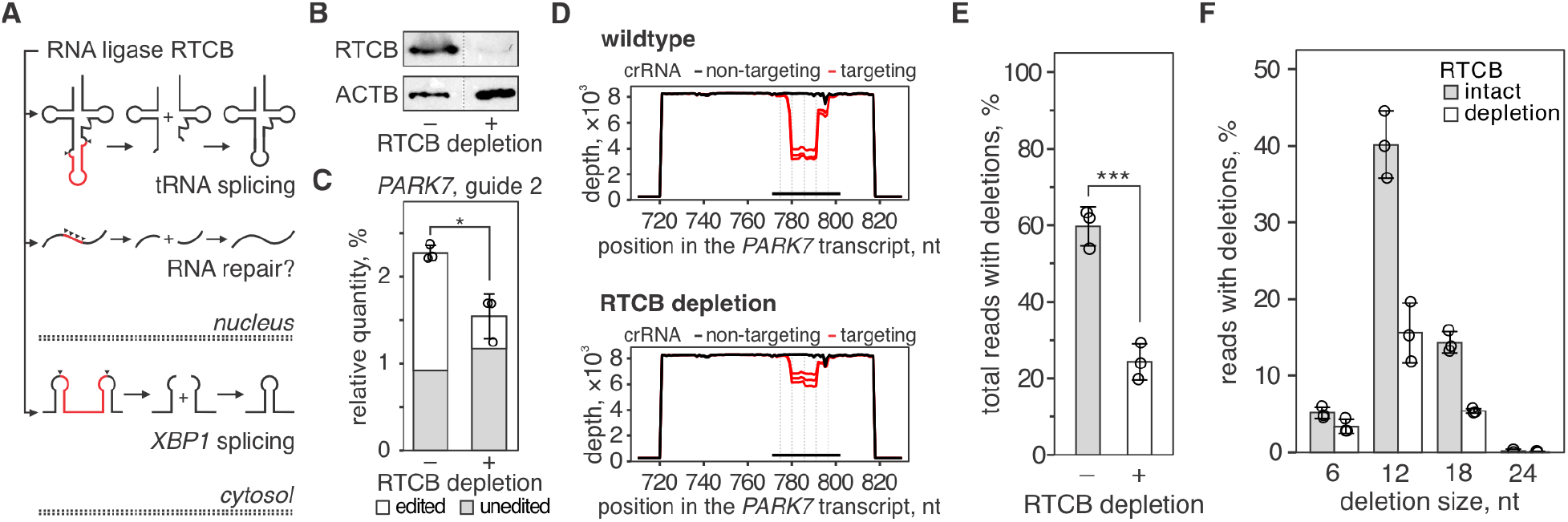
Depletion of human RNA ligase RTCB restricts the repair of type III CRISPR-mediated RNA breaks. **A**) Human RNA ligase RTCB joins RNA ends with a 2′,3′-cyclic phosphate (2’,3’>P) and a 5′-hydroxyl (5’-OH) that are produced by cellular nucleases in tRNA splicing (top) and non-canonical *XBP1* mRNA splicing (bottom) during the unfolded protein response. RNA cleavage by SthCsm generates a 2’,3’>P and a 5’-OH at each cut site, and we hypothesized that the ligase activity of RTCB is involved in the RNA repair identified in Fig. 1 (middle). **B)** Western blot with anti-RTCB or anti-ACTB (loading control) antibodies was performed with lysates of 293T cells with (+) or without (-) RTCB depletion. See the uncropped images in fig. S2. **C)** *PARK7* transcript was targeted with SthCsm (guide 2) in 293T cells with (+) or without (-RTCB depletion. *PARK7* transcript was quantified with RT-qPCR and normalized to *ACTB* non-targeting guide RNA control using the ΔΔCt method. Data are shown as the mean ± standard deviation of three biological replicates. Unequal variances t-test was used to compare mean values. *p-value < 0.05. **D)** qPCR products in (C) were sequenced, and resulting reads were aligned to the reference sequence of the *PARK7* transcript (NM_007262, GenBank). Graphs show sequencing depth (*y-axes*) at the amplified region of the transcript (*x-axes*). Every line shows a biological replicate (n = 3). The horizontal black bar indicates a region complementary to the guide RNA of the SthCsm complex. Vertical dotted lines mark predicted positions of RNA breaks. **E)** Quantification of deletions in target region of the *PARK7* transcript. Data is shown as the mean ± standard deviation of three biological replicates. Unequal variances t-test was used to compare mean values. **** p < 0.001. **F)** The distribution of different deletion outcomes in (D).

To test the effect of RTCB on RNA repair, we used SthCsm to knockdown *PARK7* transcript in cells with or without the RTCB depletion (**Fig. 2C**). In wildtype 293T cells 2.3 ± 0.1% of the target RNA was detected after transfection with SthCsm plasmid, while target RNA was ∼1.5-fold lower (1.5 ± 0.3%; p-value = 0.029) in RTCB-depleted cells. We hypothesized that the reduction of target RNA in RTCB-depleted cells is due to the decrease in RNA repair after cleavage with SthCsm. Deep-sequencing of the qPCR products spanning the target site in *PARK7* confirmed that the proportion of reads with deletions is significantly decreased in cells with depleted RTCB (**Fig. 2D**). In samples with wildtype *RTCB*, deletions of 6, 12, 18, or 24 nucleotides in target sequence are detected in 59.7 ± 5.1% of the reads, while the frequency of these deletions is 24.4 ± 4.8% (p-value < 0.001) in cells with depleted RTCB (**Fig. 2E**). Overall, the editing efficiency decreases with the depletion of RTCB, while the distribution of deletion sizes does not change (**Fig. 2F**).

### Programmable deletion of premature stop codons in RNA restores protein expression

The ClinVar database (http://www.ncbi.nlm.nih.gov/clinvar) documents more than 40,000 non-sense mutations (i.e., stop codons) in the human population that are detrimental to human health. We hypothesized that cut-and-repair RNA editing can be used to delete premature stop codons from human transcripts and restore the expression of the encoded proteins (**Fig. 3A**). To test this concept, we developed a reporter plasmid that encodes for the green fluorescent protein (GFP) fused to flag-HA epitope at the N-terminus. Between the epitope tag and the *gfp* sequence, we have inserted a stop codon to terminate the translation (stop-GFP reporter; **Fig. 3B**, top).

**Fig. 3.**
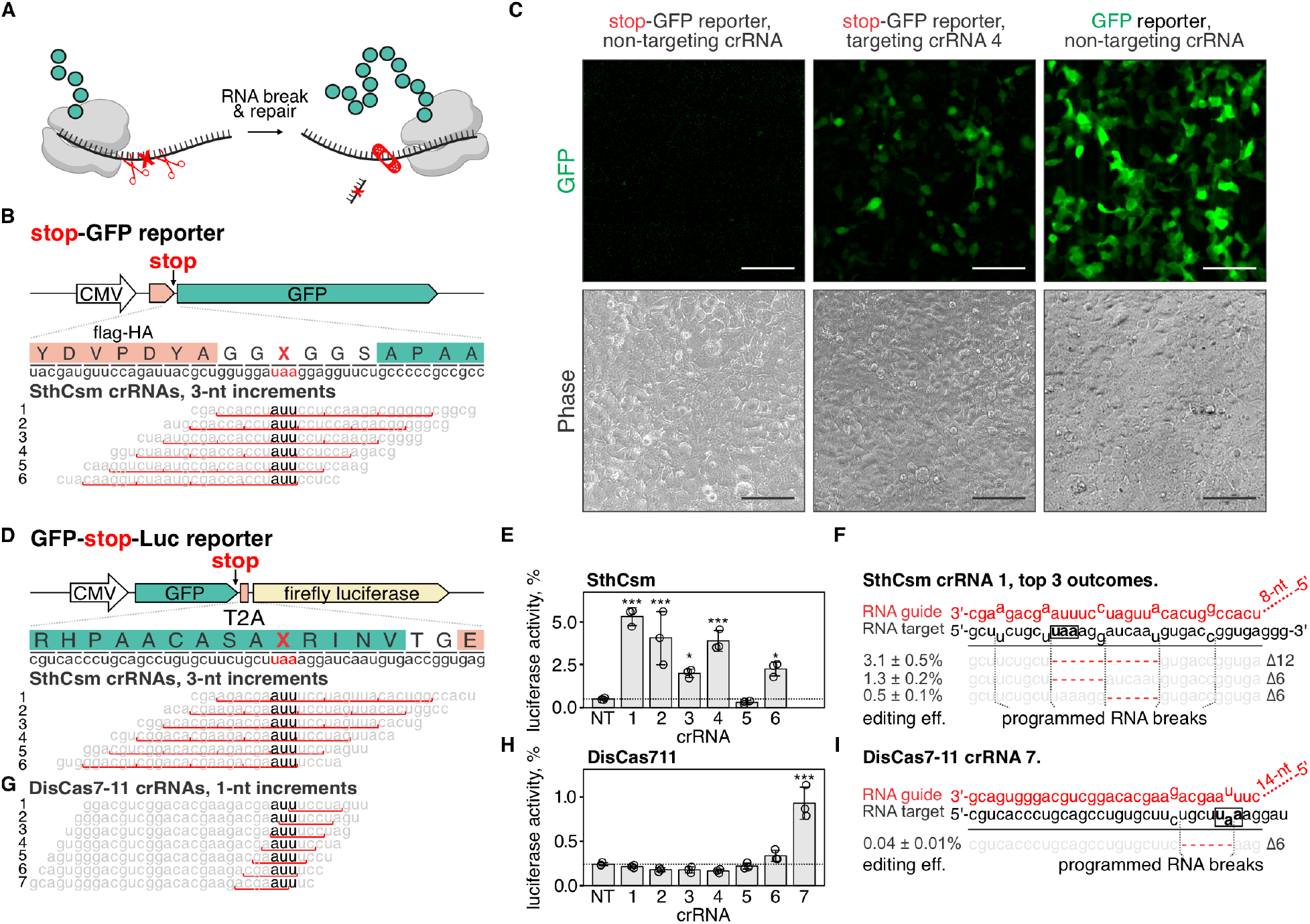
Programmable deletion of stop codons restores protein expression. **A)** Schematic representation of the proposed approach for deleting premature stop codons in human transcripts. **B)** *Top:* schematic diagram of the stop-GFP reporter plasmid. *Bottom:* Six guide RNAs for SthCsm complex were designed to excise the stop codon in the *gfp* transcript. Underlined (red) sections of target RNA are expected to be deleted. Vertical red ticks indicate predicted sites for RNA breaks. **C)** Cells were transfected with plasmids for the stop-GFP reporter and SthCsm with a non-targeting guide (left), stop-GFP reporter and SthCsm with a targeting guide (middle), or GFP reporter and SthCsm with the non-targeting guide (right). Fluorescence microscopy was used to image cells 48 h post-transfection. **D)** *Top:* schematic diagram of the GFP-stop-Luc reporter plasmid. *Bottom:* Six crRNAs for SthCsm complex were designed to delete the stop codon at the 3’-end of the *gfp* gene. **E)** Luciferase activity was measured in cell lysates 48 hours after transfection with GFP-stop-Luc and SthCsm plasmids. Luciferase activity is normalized to a control transfected with a reporter plasmid without the stop codon. Data are shown as mean ± one standard deviation of three replicates. Means were compared using one-way ANOVA, and samples with targeting guide RNAs were compared to the non-targeting control using one-tailed Dunnett’s test. * p < 0.5, ** p < 0.1, *** p < 0.001. **F)** Most frequent RNA editing outcomes in the sample with the most efficient rescue of luciferase activity in (B) (guide 1). Editing efficiency was quantified as mean ± one standard deviation of three biological replicates. Black box shows the stop codon that was targeted by type III CRISPR complexes. **G, H, I)**. The same as (D-F), but with DisCas7-11.

Transfection of the stop-GFP reporter plasmid in human cells (i.e., 293T cells) produces no detectable fluorescence (**Fig. 3C**, left). A plasmid that has the same sequence, but no stop codon, produces a robust GFP signal 48 h post-transfection (**Fig. 3C**, right). To cut out the stop codon in the reporter transcript, we designed six guides for SthCsm complex that tile across the target sequence in three-nucleotide increments (**Fig. 3B**, bottom). Transfection of the stop-GFP reporter plasmid together with three of the six SthCsm complexes (crRNA 3, 4, and 6) restores expression of the GFP protein (**Fig. 3C**, middle; **fig. S3A**).

To quantify the rescue of reporter expression, we developed a bicistronic reporter plasmid that expresses GFP and firefly luciferase proteins linked with a viral 2A peptide. At the 3’-end of the *gfp* gene, we inserted a stop codon, which permits GFP, but not luciferase expression (GFP-stop-Luc reporter; **Fig. 3D**, top). To cut out the stop codon from the reporter transcript, we designed six SthCsm guide RNAs that tile across the stop codon (**Fig. 3D**, bottom). In cells expressing SthCsm complexes that target the stop codon, we detected luciferase activity with five of the six tested crRNAs, which ranged from 2.0 ± 0.2 % with crRNA #3 to 5.3 ± 0.5% with crRNA #1 (**Fig. 3E**). Sequencing of the target RNA amplicons confirmed that samples with rescued luciferase activity contain RNA deletions that eliminate the stop codon (**Fig. 3F, fig. S3**B).

Type III-E CRISPR systems encode single polyprotein effectors that have a ribonuclease active site that processes crRNA (rather than relying of an additional nuclease, e.g., Cas6), and two ribonuclease active sites that cleave target RNA at a six-nucleotide interval (*22, 23*). These effectors are simple and compact compared to multisubunit type III complexes (i.e., SthCsm) and have the potential to produce a single editing outcome (i.e., a six-nucleotide deletion), which makes them attractive for use in RNA editing. We expressed type III-E effector from *Desulfonema ishimotonii* fused to the NLS tag (DisCas7-11) in 293T cells with seven guide RNAs tiled across the stop codon in GFP-stop-Luc reporter (**Fig. 3G**). Only crRNA #7 resulted in a statistically significant increase of the luciferase signal (0.9 ± 0.2%; p-value < 0.001), which was less efficient than SthCsm (**Fig. 3H**). In addition to editing, we quantified knockdown efficiencies of the target RNA using RT-qPCR. The RNA knockdown efficiency with SthCsm ranged from 36.8 ± 7.2 % to 54.0 ± 2.8 %, while knockdowns with DisCas7-11 were less than half as efficient (12.2 ± 5.9 % to 16.5 ± 11.2%) (**fig. S3, C and D**). Analysis of sequenced amplicons identified six-nucleotide deletions in 0.04 ± 0.01% of the reads at the site targeted with DisCas7-11 (**Fig. 3I**). Inefficient knockdown and low frequency of deletions are consistent with the inefficiency of the DisCas7-11 nuclease in biochemical assays and the low frequency of two simultaneous breaks in the target RNA (*22, 23*). While these data demonstrate the potential of type III-E effectors to produce a single RNA editing outcome, more efficient variants of Cas7-11 will be important for most applications.

## Discussion

Precise DNA manipulation, in a test tube or in a cell, has been enabled by breaking DNA at sequence-specific locations and repairing the resulting fragments. In a previous study, we applied the same concept of break-and-repair to engineer viral RNA genomes *in vitro* (*24*). Here, we extend this approach to make RNA deletions in human cells, which is facilitated by the repair of programmable RNA breaks made using type III CRISPR complexes.

We showed that the RNA ligase, RTCB is involved in the repair of RNA breaks made with type III CRISPR complexes in human transcripts. RTCB is a catalytic subunit of the tRNA ligation complex (tRLC) and is essential for tRNA splicing and unfolded protein response (*20, 25*). Repair of RNAs is understudied, and the biological significance of this process is often dismissed due to the high turnover of RNA in the cell. We have established simple GFP-based and luciferase-based reporter assays for RNA repair, which enable rapid screening of existing ligases or discovery of new RNA repair pathways. We anticipate that high-throughput assays for conditions that favor RNA repair, such as CRISPR-activation screens, will help generate a better fundamental understanding of RNA repair and ultimately lead to improved efficiency of RNA editing.

Unlike DNA editing, which often relies on protein-mediated recognition of a specific sequence motif (i.e., PAM), type III CRISPR systems only require complementary base pairing between the RNA guide and the RNA target (*15*), which eliminates additional sequence requirements and improves target site versatility. However, the frequency and distribution of RNA repair outcomes varied across different RNA targets that we tested, which may suggest that local sequence context, secondary structure of the transcript, RNA modification, or other factors may influence the efficiency of target binding, target cleavage, or repair. Screens similar to those performed for Cas13 (*26, 27*) will help establish predictive models that enhance the efficiencies of RNA editing and help advance this technology for therapeutic applications.

## Materials and Methods

### Plasmids

pDAC627 expressing type III CRISPR-associated proteins from *Streptococcus thermophilus* (Csm2 – Csm5, Cas10, and Cas6) fused to a nuclear localization signal (NLS) and RNA guide was a gift from Jennifer Doudna (Addgene plasmid #195240; http://n2t.net/addgene:195240; RRID: Addgene_195240) (*16*). To clone SthCsm guides targeting *PPIB, PARK7*, or reporter transcripts (stop-GFP and GFP-stop-Luc), pairs of complementary DNA oligos (table S1) were annealed and phosphorylated with T4 polynucleotide kinase [New England Biolabs (NEB)] as previously described (*3*). The resulting fragments were cloned using the *Bbs*I restriction enzyme (NEB) into pDAC627. The unmodified pDAC627 vector was used as the non-targeting control. Guide RNAs targeting *PPIB* and *PARK7* transcripts were designed using the *cas13design* algorithm (https://gitlab.com/sanjanalab/cas13), which was customized to select 32-nucleotide-long guide RNAs (*26*). Highest-ranking guide RNAs that target different regions of the transcripts were selected.

pSpCas9(BB)-2A-Puro (PX459) was a gift from Feng Zhang (Addgene plasmid #48139; http://n2t.net/addgene:48139;RRID:Addgene_48139). Cas9 sgRNAs targeting the *RTCB* gene were cloned as previously described (table S2) (*3*). Guide sequences were selected in second and third exons of the gene using inDelphi web interface based on predicted knockout efficiency and editing precision (https://indelphi.giffordlab.mit.edu/) (*28*).

pDF0159 pCMV - huDisCas7-11 mammalian expression plasmid was a gift from Omar Abudayyeh & Jonathan Gootenberg (Addgene plasmid #172507; http://n2t.net/addgene:172507; RRID:Addgene_172507) (*18*). To add the NLS tag to the N-terminus of the DisCas-11 protein, the plasmid pDF0159 was modified using Q5 site-directed mutagenesis (table S3).

pDF0114 pu6-eco31i-eco31i-dis7-11-mature-dr-guide-scaffold was a gift from Omar Abudayyeh & Jonathan Gootenberg (Addgene plasmid #186981; http://n2t.net/addgene:186981 ; RRID: Addgene_186981) (*29*). To clone DisCas7-11 guides, two partially complementary DNA oligos were annealed and phosphorylated (table S4). The resulting duplexes were cloned into pDF0114 using *Bsa*I restriction enzyme (NEB) to substitute a non-targeting guide sequence.

To generate reporter plasmids pStop-GFP and pGFP-stop-Luc, dscGFP-2A-Luciferase gene cassette was amplified from pGreenFire1-ISRE (EF1α-puro) vector (System Biosciences, TR016PA-P) using PCR and cloned in the pEGFP-N1 vector (Clontech) at the *Kpn*I and *Not*I restriction sites. Restriction sites and Flag-HA tag at the N-terminus of the GFP were added into the overhangs of the oligonucleotide primers that were used for PCR. Stop codons were inserted in the 5’-or the 3’-end of the *gfp* gene using Q5 site-directed mutagenesis.

All plasmid sequences were confirmed with whole-plasmid sequencing at Plasmidsaurus (http://www.plasmidsaurus.com/).

### Nucleic acids

All DNA oligos were purchased from Eurofins (table S1-S6).

### Antibodies

Antibodies used for western blot: rabbit anti-RTCB (Proteintech, Cat:19809-1-AP, 1:1000), rabbit anti-ACTB (ABclonal, Cat: AC026, 1:20,000), and goat anti-rabbit IgG peroxidase-conjugated (Jackson ImmunoResearch, Cat: 111-035-003, 1:10,000).

### Cell cultures

293T cells (ATCC, CRL-3216) were maintained at 37°C and 5% CO2 in Dulbecco’s Modified Eagle Medium (DMEM, GIBCO, cat. #12100-061) supplemented with 10% fetal bovine serum (FBS, ATLAS Biologicals, Lot. #F31E18D1), sodium bicarbonate (3.7 g/L), 50 I.U./mL penicillin and 50 mg/mL streptomycin. The cells tested PCR-negative for mycoplasma.

### RTCB depletion

To deplete RTCB, 293T cells were transfected using Lipofectamine 3000 (ThermoFisher Scientific) with the plasmid encoding for Cas9, puromycin resistance protein, and a guide RNA targeting the *RTCB* gene. Twenty-four hours post-transfection, the media was changed to the media with puromycin (1 µg/mL), and the cells were selected for three days. After selection, cells were seeded on 96-well plates at 3 cells per well density to generate clonal cell lines. Cell clones were monitored for 14 days, and wells with single colonies were marked and used to determine knockout efficiencies. To determine knock-out efficiency, the genomic DNA from bulk populations or clonal cell lines was extracted using DNeasy Blood & Tissue Kit (QIAGEN) and targeted genome regions were amplified with Q5 polymerase (NEB) (table S5). Amplicons were Sanger-sequenced (Psomagen), and resulting traces (i.e., ab1 files) were used to quantify indels with the ICE software from Synthego (https://ice.synthego.com) (*30*).

### Plasmid transfection

293T cells were seeded one or two days before the experiment and transfected when ∼80-90% confluency was reached. For the experiment shown in Fig. 1B and Fig. 2C, 500 ng of plasmid DNA encoding for SthCsm-complex and guide RNA was mixed with 1.5 *μ*L of FuGENE HD reagent (Promega), incubated for 15 min, and added to the well of a 24-well plate. Twelve hours post-transfection, the cells were detached and seeded on 12-well plates in a media with puromycin (1 µg/mL). Three days after puromycin selection, the total RNA was extracted from cells using RNeasy Plus Mini Kit (QIAGEN, # 74134).

For the experiment shown in Fig. 3, the cells were seeded in a 96-well plate in ∼80-90% confluency. For editing with SthCsm-complex, 100 ng of plasmid encoding for SthCsm and 50 ng of plasmid encoding for a reporter (stop-GFP or GFP-stop-Luc) were mixed with 0.8 *μ*L FuGENE HD reagent (Promega), incubated for 10 min, and added to the well. For editing with DisCas7-11, 100 ng of plasmid encoding for DisCas7-11, 50 ng of the plasmid encoding for guide RNA, and 50 ng of the plasmid encoding for the GFP-stop-Luc reporter were mixed with 1.1 *μ* FuGENE HD reagent (Promega), incubated for 10 min, and added to the wells. Forty-eight hours post-transfection, the cells were imaged with the Nikon Ti-Eclipse inverted microscope (Nikon Instruments) equipped with a SpectraX LED excitation module (Lumencor) and emission filter wheels (Prior Scientific) (for the reporter stop-GFP). Fluorescence imaging used excitation/emission filters and dichroic mirrors for GFP (Chroma Technology Corp.). Images were acquired with Plan Fluor 20Ph objective and an iXon 896 EM-CCD camera (Andor

Technology Ltd.) in NIS-Elements software. For luciferase assays, media was removed, and cells were lysed in 50 µL of the 1X Cell Culture Lysis Reagent (Promega). Lysates were used for luciferase assays (20 µL) with Luciferase assay system (Promega) and RNA extraction (20 µL) with RNeasy Plus Mini Kit (QIAGEN, # 74134).

### RT-qPCR

Total RNA from cells was extracted with RNeasy Plus Mini Kit (QIAGEN, # 74134). The reverse-transcription (RT) reactions were performed with 2X LunaScript RT SuperMix Kit (NEB) with 5 *μ*L of RNA input (250 ng of RNA total) in 10 *μ*L total reaction volume. The RT reactions containing cDNA were diluted 100-fold, and 5 µL was used for qPCR reactions with 2X Universal SYBR Green Fast qPCR Mix (ABClonal). Each qPCR reaction contained 5 µL of cDNA, 4.2 µL of Nuclease-free Water, 0.4 µL of 10 mM Primers (table S6), and 10 µL of 2X mastermix. Nuclease-free water was used as no template control (NTC). Three technical replicates were performed for each sample. Amplification was performed in QuantStudio 3 Real-Time PCR System instrument (Applied Biosystems) as follows: 95°C for 3 min, 40 cycles of 95°C for 5 s, and 60°C for 30 s, followed by Melt Curve analysis. Results were analyzed in Design and Analysis application at Thermo Fisher Connect Platform.

### Nanopore sequencing

Amplicons from RT-qPCR reactions were purified with magnetic beads (Mag-Bind® TotalPure NGS), and ∼13 ng of DNA was used to prepare sequencing libraries as described in SQK-LSK114 protocol using Native Barcoding Kit 24 V14 (SQK-NBD114.24). ∼20 fM of the library was loaded on the Nanopore MinION [R10.4.1 flow cell, accurate mode (260 bases per second)]. The flow cell was primed, and sequencing libraries were loaded according to the Oxford Nanopore protocol (SQK-LSK114). The sequencing run was performed in the fast base calling mode in the MinKNOW software. After the run, raw sequencing data (POD5 files) was re-basecalled with the super-accuracy model (dna_r10.4.1_e8.2_260bps_sup@v4.1.0) using Dorado basecaller (v0.3.3) and demultiplexed with guppy_barcoder (both from Oxford Nanopore).

### Sequencing data analysis

Sequencing reads were aligned to the reference using *minimap2* (v2.17 -r954-dirty) with Nanopore preset (-ax map-ont setting). Alignments were converted to BAM format, sorted, and indexed using *samtools* v1.13. Sequencing reads were trimmed to remove primer binding regions using the *samtools ampliconclip*. To make sequencing depth plots, a subset of reads was randomly selected using *samtools view*. Sequencing depth was computed with the *samtools depth*. Deletions in sequencing reads were quantified using the extract-junctions.py script (github.com/hyeshik/sars-cov-2-transcriptome/blob/master/nanopore/_scripts/extract-junctions.py) (*31*). Deletion counts at the SthCsm target sites were normalized to the number of reads that span the target region, which was computed by extracting read information from BAM files using the *bamtobed* function in *bedtools* v2.30.0 package and filtering reads by start and end coordinate (Data S1). All plots were created using *ggplot2* v3.3.5 package in RStudio software v2023.03.0.

### Western blot

Cells were washed two times with PBS and lysed in RIPA buffer (150 mM sodium chloride, 1% NP-40, 0.5% sodium deoxycholate, 0.2% SDS, 50 mM Tris, pH 8.0) at 4°C for 30 min. The lysates were clarified by centrifugation (20 min at 10,000 g) and stored at -80°C. Lysates were mixed with 6X Laemmli buffer, heated at 98°C for 5 min, resolved in 12% SDS-PAGE gel, and transferred onto a nitrocellulose membrane using Mini Trans-Blot Electrophoretic Transfer Cell (Bio-Rad, #1703930). Membranes were stained with Ponceau S, imaged, washed, blocked, and probed with indicated antibodies. The proteins were visualized using Pierce ECL Western Blotting Substrate (ThermoFisher Scientific, #32106) and exposed to X-Ray film (sc-201696, Santa Cruz Biotech).

### Statistical analysis

All experiments were performed in triplicates. All RT-qPCR reactions were performed in three technical replicates. Statistical comparisons in Fig. 2 were performed in R with Welch’s unequal variances *t*-test using the *t*.*test* function from the *stats* package. One-way ANOVA (Fig. 3) was performed using the *aov* function from the *stats* package in R. Post-hoc pairwise comparisons were performed with one-tailed Dunnett’s test using the *glht* function from the *multcomp* package in R. Significance levels: \**p* < 0.05, *\**\**p* < 0.01, \*\*\**p* < 0.001.

## Acknowledgments

Base-calling of Nanopore sequencing data was performed with the Tempest High Performance Computing System, which is maintained by University Information Technology Research Cyberinfrastructure at Montana State University. We thank Dr. Matthew Taylor for the generous use of the fluorescent microscope. Some diagrams and schematics were created with BioRender.com.

## Author contributions

Conceptualization: A. Nemudryi, A. Nemudraia, and B.W.

Methodology: A. Nemudryi and A. Nemudraia.

Investigation: A. Nemudryi and A. Nemudraia.

Formal analysis: A. Nemudryi and A. Nemudraia.

Visualization: A. Nemudryi and A. Nemudraia.

Supervision: A. Nemudryi, A. Nemudraia, and B.W.

Funding acquisition: B.W. and A. Nemudryi.

Writing - Original Draft: A. Nemudryi and A. Nemudraia.

Writing - Review & Editing: B.W., A. Nemudryi, and A. Nemudraia.

## Data and materials availability

Raw sequencing data are deposited to NCBI Sequence Read Archive (SRA) under a single Bioproject accession ID. Data will be released upon publication. Code for data analysis will be deposited to Zenodo upon publication. Plasmids are available from the corresponding author upon request.

