## Supplemental Figures and Tables for "Repair of CRISPR-guided RNA breaks enables site-specific RNA editing in human cells"

**The PDF file includes:**

Figs. S1 to S3
Tables S1 to S6

**Other Supplementary Materials for this manuscript include the following:**

Data S1 (separate file)

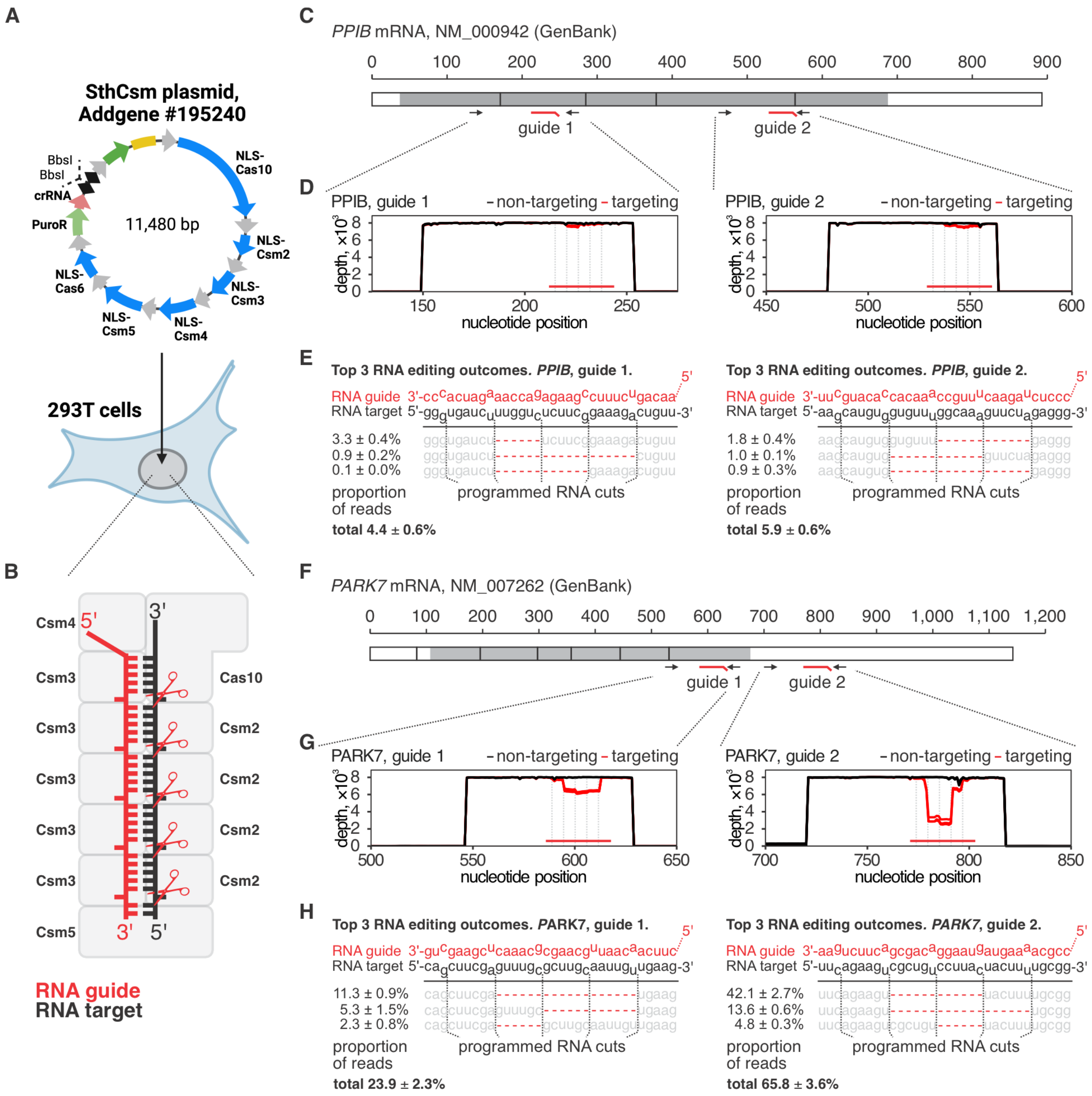


**Fig. S1. The distribution and frequency of RNA editing outcomes in *PPIB* and *PARK7* mRNAs. A)** Diagram of the plasmid (Addgene #195240) encoding for human codon-optimized and NLS-tagged protein subunits of the type III-A CRISPR-Csm complex from *Streptococcus thermophilus* (SthCsm), CRISPR RNA (crRNA) -processing nuclease Cas6, and single-spacer CRISPR array. CRISPR repeats are shown as black diamonds. The sequence between the repeats has two BbsI restriction sites for cloning spacers for targeting complementary target RNAs. **B)** Diagram of the assembled SthCsm ribonucleoprotein complex bound to the target RNA (black). crRNA is shown with red color. The target RNA is cleaved in six nucleotide intervals by Csm3 nucleases in the backbone of the complex. Cleavage sites are indicated with red scissors. **C)** Diagram of the *PPIB* mRNA (NM_000942, GenBank). The scale shows nucleotide position along the transcript. Blocks show exons in the spliced mRNA, gray color indicates the position of the PPIB open reading frame (ORF). Red lines mark the position of the SthCsm guide RNAs.

Black arrows indicate position of the binding sites for oligonucleotide primers used for RT-qPCR in Fig. 1B. **D)** qPCR amplicons were deep-sequenced, and resulting reads were aligned to the reference sequences. A subset of 8,000 reads was randomly selected from the alignment for the representation. Plots show sequencing coverage (y-axes) along the length of the amplicons (x-axes). Each line represents a single replicate, three replicates total per guide RNA. Vertical dotted gray lines indicate predicted RNA break sites. The horizontal red line shows the position of the guide RNA with respect to the target. **E)** Top three most frequent RNA editing outcomes in PPIB transcript. Dotted lines indicate the positions of RNA breaks introduced with the CRISPR complex. Red dashes depict deletions identified in the sequencing data. Deletion frequency was quantified as the mean ± one standard deviation of three biological replicates. **F)** Same as in (**C**) but for the *PARK7* transcript (NM_007262, GenBank). **G)** Same as in (**D**) but for target sites in the *PARK7* transcript. **H)** Top three most frequent RNA editing outcomes at the target sites in *PARK7*. Data is shown as the mean ± one standard deviation of three biological replicates.

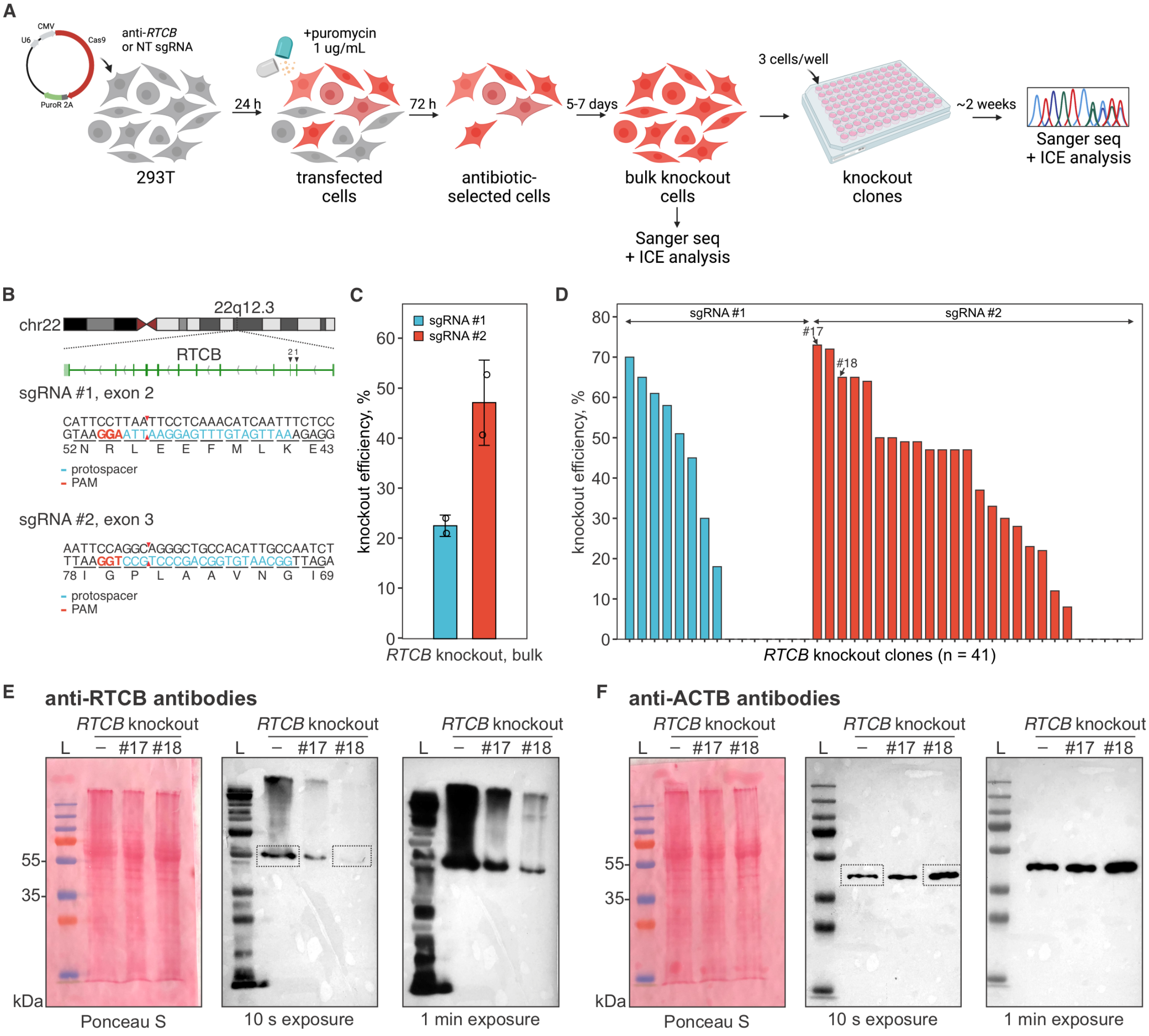


**Fig. S2. Knockout of RTCB gene with CRISPR-Cas9. A)** Schematics of the knockout strategy. **B)** Diagram showing the exon-intron structure of the *RTCB* gene (top) and sequences targeted with Cas9 (middle and bottom). **C)** *RTCB* knockout efficiency in bulk cells was quantified by analyzing Sanger traces from target site amplicons using the ICE web tool (Synthego). **D)** Quantification of *RTCB* knockout efficiency in 293T cell clones. **E and F)** Clones #17 and #18, indicated with black arrows in (**D**), were lysed, and lysates were probed with anti-RTCB or anti-ACTB antibodies. Dotted boxes show parts of the images that were cropped out for main Fig. 2B.

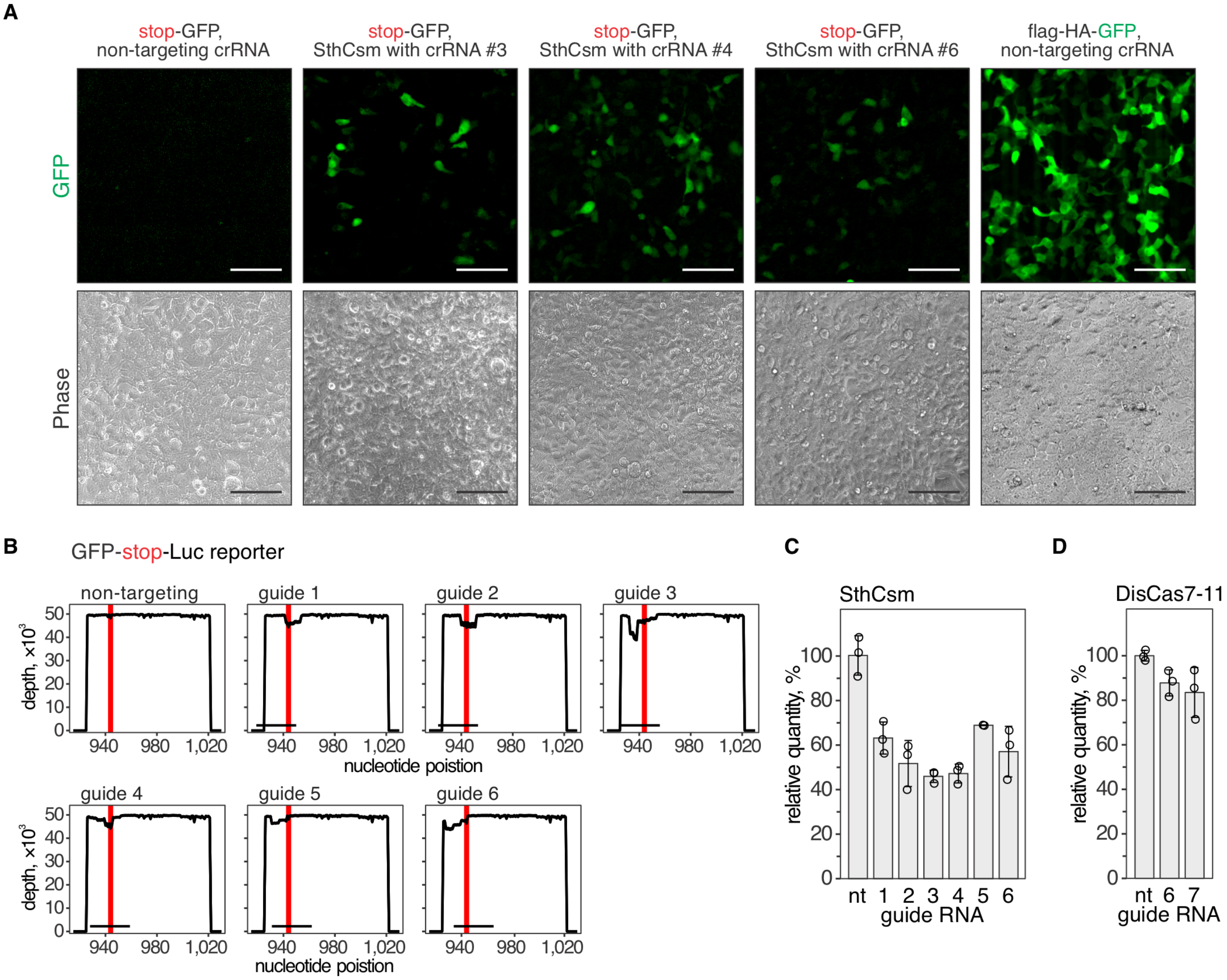


**Fig. S3. Programmable deletion of stop codons in RNA restores protein expression.**

**A)** Fluorescent imaging of cells transfected with plasmids for the stop-GFP reporter and SthCsm with a non-targeting guide (left), stop-GFP reporter and SthCsm with targeting guides, or GFP reporter and SthCsm with the non-targeting guide (right). **B)** Deep-sequencing of qPCR amplicons in (C). Reads were aligned to the reference sequence of the GFP-stop-Luc reporter. Graphs show sequencing depth (y-axes) at the amplified region of the transcript (x-axes). Every line shows a biological replicate (n = 3). The horizontal black bar indicates a region complementary to the guide RNA of the SthCsm complex. Vertical red line marks the postion of the stop codon targeted with SthCsm. **C)** RNA was extracted from the same lysates that were used in luciferase assays shown in Fig. 3E. RT-qPCR was performed with oligonucleotide primers that flank the region targeted with SthCsm in the GFP-stop-Luc reporter transcript. Transcript quantities are normalized to *ACTB* and non-targeting guide RNA control using the ΔΔCt method. The mean ± one standard deviation of three biological replicates is shown. nt – non-targeting guide RNA. **D)** Same as in (C), but for the experiment with DisCas7-11 in Fig. 3H.

**Table S1.**

Oligonucleotides used for cloning SthCsm guide RNAs

| Name | Sequence 5`-3` |
|---|---|
| *Guides targeting endogenous transcripts* | |
| PPIB_g1F | AAACAACAGTCTTTCCGAAGAGACCAAAGATCACCC |
| PPIB_g1R | TATCGGGTGATCTTTGGTCTCTTCGGAAAGACTGTT |
| PPIB_g2F | AAACCCCTCTAGAACTTTGCCAAACACCACATGCTT |
| PPIB_g2R | TATCAAGCATGTGGTGTTTGGCAAAGTTCTAGAGGG |
| PARK7_g1F | AAACCTTCAACAATTGCAAGCGCAAACTCGAAGCTG |
| PARK7_g1R | TATCCAGCTTCGAGTTTGCGCTTGCAATTGTTGAAG |
| PARK7_g2F | AAACCCGCAAAAGTAGTAAGGACAGCGACTTCTGAA |
| PARK7_g2R | TATCTTCAGAAGTCGCTGTCCTTACTACTTTTGCGG |
| *Guides targeting stop-GFP reporter* | |
| sthcsm_stop_gfp_g1F | AAACGCGGCGGGGGCAGAACCTCCTTATCCACCAGC |
| sthcsm_stop_gfp_g1R | TATCGCTGGTGGATAAGGAGGTTCTGCCCCCGCCGC |
| sthcsm_stop_gfp_g2F | AAACGCGGGGGCAGAACCTCCTTATCCACCAGCGTA |
| sthcsm_stop_gfp_g2R | TATCTACGCTGGTGGATAAGGAGGTTCTGCCCCCGC |
| sthcsm_stop_gfp_g3F | AAACGGGGCAGAACCTCCTTATCCACCAGCGTAATC |
| sthcsm_stop_gfp_g3R | TATCGATTACGCTGGTGGATAAGGAGGTTCTGCCCC |
| sthcsm_stop_gfp_g4F | AAACGCAGAACCTCCTTATCCACCAGCGTAATCTGG |
| sthcsm_stop_gfp_g4R | TATCCCAGATTACGCTGGTGGATAAGGAGGTTCTGC |
| sthcsm_stop_gfp_g5F | AAACGAACCTCCTTATCCACCAGCGTAATCTGGAAC |
| sthcsm_stop_gfp_g5R | TATCGTTCCAGATTACGCTGGTGGATAAGGAGGTTC |
| sthcsm_stop_gfp_g6F | AAACCCTCCTTATCCACCAGCGTAATCTGGAACATC |
| sthcsm_stop_gfp_g6R | TATCGATGTTCCAGATTACGCTGGTGGATAAGGAGG |
| *Guides targeting GFP-stop-Luc reporter* | |
| csm_gfp_stop_luc_g1F | AAACTCACCGGTCACATTGATCCTTTAAGCAGAAGC |
| csm_gfp_stop_luc_g1R | TATCGCTTCTGCTTAAAGGATCAATGTGACCGGTGA |
| csm_gfp_stop_luc_g2F | AAACCCGGTCACATTGATCCTTTAAGCAGAAGCACA |
| csm_gfp_stop_luc_g2R | TATCTGTGCTTCTGCTTAAAGGATCAATGTGACCGG |
| csm_gfp_stop_luc_g3F | AAACGTCACATTGATCCTTTAAGCAGAAGCACAGGC |
| csm_gfp_stop_luc_g3R | TATCGCCTGTGCTTCTGCTTAAAGGATCAATGTGAC |
| csm_gfp_stop_luc_g4F | AAACACATTGATCCTTTAAGCAGAAGCACAGGCTGC |
| csm_gfp_stop_luc_g4R | TATCGCAGCCTGTGCTTCTGCTTAAAGGATCAATGT |
| csm_gfp_stop_luc_g5F | AAACTTGATCCTTTAAGCAGAAGCACAGGCTGCAGG |
| csm_gfp_stop_luc_g5R | TATCCCTGCAGCCTGTGCTTCTGCTTAAAGGATCAA |
| csm_gfp_stop_luc_g6F | AAACATCCTTTAAGCAGAAGCACAGGCTGCAGGGTG |
| csm_gfp_stop_luc_g6R | TATCCACCCTGCAGCCTGTGCTTCTGCTTAAAGGAT |

**Table S2.**

Oligonucleotides used for cloning Cas9 guides targeting RTCB gene

| Name | Sequence 5`-3` |
|---|---|
| RTCB_sg1F | CACCGAATTGATGTTTGAGGAATTA |
| RTCB_sg1R | CAAATAATTCCTCAAACATCAATTC |
| RTCB_sg2F | CACCGGCAATGTGGCAGCCCTGCC |
| RTCB_sg2R | CAAAGGCAGGGCTGCCACATTGCC |

**Table S3.**

Oligonucleotide primers used for Q5 SDM

| Name | Sequence 5`-3` |
|---|---|
| NLS-DisCas711_F | AAGCGGAAGGTCGGCGGTAGCACTACTATGAAGATTTCAATTGAATTC |
| NLS-DisCas711_R | CTTCTTTGGGGCCATGGTGGCGGCTCTCCCTATAGTG |

**Table S4.**

Oligonucleotides used for cloning DisCas7-11 guide RNAs

| Name | Sequence 5`-3` |
|---|---|
| discas711_gfp_stop_luc_g1F | GAACTTGATCCTTTAAGCAGAAGCACAGGCTGCAGG |
| discas711_gfp_stop_luc_g1R | AAAACCTGCAGCCTGTGCTTCTGCTTAAAGGATCAA |
| discas711_gfp_stop_luc_g2F | GAACTGATCCTTTAAGCAGAAGCACAGGCTGCAGGG |
| discas711_gfp_stop_luc_g2R | AAAACCCTGCAGCCTGTGCTTCTGCTTAAAGGATCA |
| discas711_gfp_stop_luc_g3F | GAACGATCCTTTAAGCAGAAGCACAGGCTGCAGGGT |
| discas711_gfp_stop_luc_g3R | AAAAACCCTGCAGCCTGTGCTTCTGCTTAAAGGATC |
| discas711_gfp_stop_luc_g4F | GAACATCCTTTAAGCAGAAGCACAGGCTGCAGGGTG |
| discas711_gfp_stop_luc_g4R | AAAACACCCTGCAGCCTGTGCTTCTGCTTAAAGGAT |
| discas711_gfp_stop_luc_g5F | GAACTCCTTTAAGCAGAAGCACAGGCTGCAGGGTGA |
| discas711_gfp_stop_luc_g5R | AAAATCACCCTGCAGCCTGTGCTTCTGCTTAAAGGA |
| discas711_gfp_stop_luc_g6F | GAACCCTTTAAGCAGAAGCACAGGCTGCAGGGTGAC |
| discas711_gfp_stop_luc_g6R | AAAAGTCACCCTGCAGCCTGTGCTTCTGCTTAAAGG |
| discas711_gfp_stop_luc_g7F | GAACCTTTAAGCAGAAGCACAGGCTGCAGGGTGACG |
| discas711_gfp_stop_luc_g7R | AAAACGTCACCCTGCAGCCTGTGCTTCTGCTTAAAG |

**Table S5.**

Oligos used to amplify Cas9-targeted regions in the *RTCB* gene

| Name | Sequence 5`-3` |
|---|---|
| RTCB_seq_F1 | AGATGGAGCTTCAAATCCGTT |
| RTCB_seq_R1 | TGTGGTTTTGAGGGTATGAGAAT |
| RTCB_seq_F2 | GGTTATGTCTGGCTGTCCAAAG |
| RTCB_seq_R2 | ATCTGGAATTTGAAGCTGGGTAG |

**Table S6.**

Primers used for RT-qPCR

| Name | Sequence 5`-3` |
|---|---|
| ACTB_F | AGAGCCTCGCCTTTGCC |
| ACTB_R | ATGCCGGAGCCGTTGTC |
| PPIB_cr1_qF1 | GCGGCCGATGAGAAGAAGAA |
| PPIB_cr1_qR1 | AGCTAAGGCCACAAAATTATCCAC |
| PPIB_cr2_qF1 | CGCAGGCAAAGACACCAAC |
| PPIB_cr2_qR1 | TTCCGCACCACCTCCA |
| PARK7_cr1_qF1 | CCTACTCTGAGAATCGTGTGGA |
| PARK7_cr1_qR1 | GAGCCGCCACCTCCTTG |
| PARK7_cr2_qF1 | AGAGAAACAGGCCGTTAGGA |
| PARK7_cr2_qR1 | GGCTGAGAAATCTCTGTGTAGTTG |
| gfp_stop_luc_qF1 | GGATGGACCGTCACCCTG |
| gfp_stop_luc_qR1 | TGTTTTTGGCGTCTTCCATACC |

**Data S1. (separate file)**
Analysis of the sequencing data in Figs. 1-3, fig. S1 and fig. S3
